# PHOSPHATE-INDUCED 1 and its paralogs positively regulate cellulose biosynthesis in Arabidopsis

**DOI:** 10.1101/2024.01.24.577082

**Authors:** Fugen Yu, Qiang Wang, Jieren Qiu, Yonghui Liao, Xin Wang, Tianjun Cao, Jiao Zhang, Shan Yin, Zhong Zhuang, Xiaolin Chen, Haojie Jiang, Shan Lu

## Abstract

Cellulose is not only the most abundant biopolymer on Earth but also an essential multi-purpose resource, primarily synthesized by cellulose synthase complexes (CSCs) located on plasma membrane in vascular plants. Although insights into the functions of CSCs have been provided by the identification of cellulose synthase catalytic subunits, our understanding of the regulation of CSC activity remains limited. In this study, combining molecular and genetic methods, we demonstrated that both PHI-1 and its paralogs residing in the plasma membrane are involved in the cellulose biosynthesis regulation as a positive regulatory component of CSCs by feeding developmental and environmental cues into their activities in Arabidopsis (*Arabidopsis thaliana*). Our finding reveals a mechanism for how plants regulate cellulose synthesis in response to developmental and environmental signals, and thus improves the likelihood that cellulose biosynthesis could be feasibly adapted for sustainable purposes.

## Introduction

Cellulose is the most abundant biopolymer on Earth. As the major load-bearing polymer in plant cell walls, cellulose not only constructs the plant body morphology, provides the plant with mechanical support, and contributes to plant resistance to biotic and abiotic stresses, but is also used as raw materials for industries (Cosgrove, 2005; Somerville, 2006; Endler and Persson, 2011). More recently, cellulose has drawn the attention of the public as a promising renewable biofuel source to combat global climate change for sustainable development (Burton and Fincher, 2014; Yoshida et al., 2021). Generally, it is believed that cellulose is synthesized by protein complexes known as cellulose synthase complexes (CSCs) located at the plasma membrane and presenting a rosette shape although their precise stoichiometry is unclear (Giddings et al., 1980; Mueller and Brown, 1980; Arioli et al., 1998; McFarlane et al., 2014). The cellulose synthase catalytic subunits (CESAs) were well-known as the catalytic components of CSCs and there are ten CESA paralogs in Arabidopsis (Somerville, 2006; Endler and Persson, 2011; McFarlane et al., 2014). Of which, CESA1, CESA3 and CESA6-likes are involved in the formation of the primary cell wall while CESA4, CESA7 and CESA8 are involved in the formation of the secondary cell wall (Fagard et al., 2000; Taylor et al., 2000; Scheible et al., 2001; Desprez et al. 2002; Ellis et al., 2002; Desprez et al., 2007; Persson et al., 2007). In addition, several proteins were also demonstrated to have a direct connection with the CSCs (Li et al., 2012; Vain et al., 2014; Endler et al., 2015).

However, the mechanisms responsible for regulating the activity of the CSCs are still poorly understood, especially in defining the regulatory components that specifically impact the CSC activity. The plant cell wall is a dynamic structure that changes in response to developmental and environmental cues, yet there is little evidence for environmental regulation of *CESA* expression (Somerville, 2006).

Therefore, it is believed that the activity of CSC is not entirely regulated at the transcriptional level of *CESA*s, but at the post-transcriptional level of *CESA*s to a certain degree in plants (Somerville, 2006), just as in bacteria (Römling and Galperin, 2015). One of the ways is to regulate the activity of CSC by a protein located on the plasma membrane interacting with CESAs in response to developmental and environmental signals (Somerville, 2006; Endler and Persson 2011). Because of the redundancy of gene function, it is difficult to identify the regulatory components of the CSC using the reverse genetics strategy (Somerville, 2006; Pedersen et al., 2023). So far, no positive regulatory component of CSC has been identified.

When we finished the research project on the coordinated control mechanism between cell cycle activities and cell morphogenesis, we happened to meet a transgenic plant line whose T-DNA was inserted into the promoter region of *PHOSPHATE-INDUCED 1* (*PHI-1*, also known as *EXORDIUM LIKE 1*, encoded by *AT1G35140*) (https://www.arabidopsis.org/servlets/TairObject?id=137481&type=locus). By searching literature, it was found that its homologous gene was originally identified in a synchronization experiment of cell division of tobacco cultured cells (Toshio et al., 1999), while its closest paralogous gene *EXORDIUM* (*EXO*) expressed in meristem in a promoter trapping experiment in Arabidopsis (Farrar et al., 2003). These experimental results seemed to indicate that PHI-1 is related to cell cycle activities, and our observation of this transgenic plant line showed that it was a little premature. Hence, we are interested in further research on it.

In this study, we found that genes directly connected with *PHI-1* on the co-expressed gene network are mainly cell wall-related and *PHI-1* responds to the well-known developmental and environmental cues related to cellulose synthesis. Our experimental results showed that the cell-type-specific expression pattern of *PHI-1* is consistent with a need for cellulose synthesis in various cells, and overexpression of *PHI-1* raised the cellulose content, decreased the cell wall plasticity and resulted in a distinct alteration of plant architecture as well. We also found that the loss-of-function of the entire PHI-1 family weakened cell anisotropy resulting from the cellulose reduction. In addition, we identified both PHI-1 and its paralogs as plasma membrane proteins and PHI-1 can interact directly with multiple CESA members and overexpression of *PHI-1* can rescue *cesa1*mutant. Taken together, we present a mechanism that the PHI-1 family protein as a positive regulatory component of the CSC adjusts dynamically cellulose synthesis in response to developmental and environmental signals.

## Results

### PHI-1 is a potential regulating hub of cellulose biosynthesis

In Arabidopsis, PHI-1 is a protein of 309 amino acid residues, containing the PHI-1 conserved region and belongs to a protein family of highly conserved land plant-specific proteins (Phosphate-responsive 1 family protein) (https://www.ncbi.nlm.nih.gov/gene/840399) (Farrar et al., 2003; Schröder et al., 2009) (Figure 1A). The mRNA of its homologous gene was first isolated in studying cell cycle synchronization of tobacco BY-2 cells when it is re-activated in cultured tobacco cells following release from phosphate starvation-induced cell cycle arrest (Toshio et al., 1999). In Arabidopsis, *EXO* is *PHI-1*’s closest paralogous gene (Figure 1A), which was expressed in proliferating cells and played a role in meristem function, first identified by promoter trapping (Farrar et al., 2003). However, information from the public database ATTED-II showed that genes directly connected with *PHI-1* on the co-expressed gene network are mainly cell wall-related (Obayashi et al., 2022) (Figure 1B), and some experiments indicated that *PHI-1* was involved in carbon balance in plant (Baena-González et al., 2007; Schröder et al. 2011).

**Figure 1.**
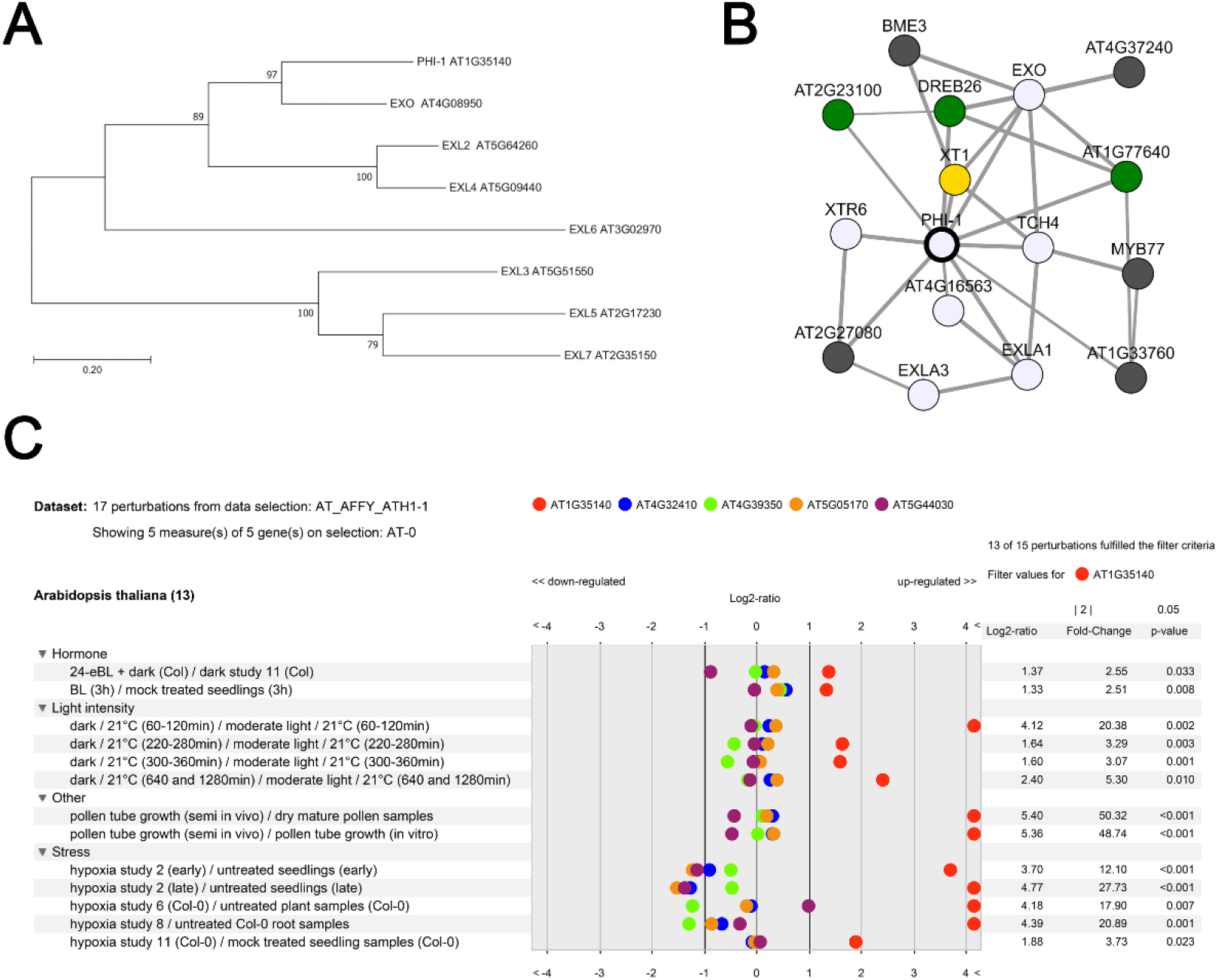
PHI-1 is a potential regulating hub of cellulose biosynthesis. (A) Molecular phylogenetic analysis of PHI-1 protein family in Arabidopsis. The tree is drawn to scale, with branch lengths measured in the number of substitutions per site. On the scale, 0.2 represents the estimated number of substitutions per site. (B) Genes directly connected with *PHI-1* on the co-expressed gene network. Of which, *TCH4*, *XT1*, *XTR6* and *EXLA1* are well-known cell wall-related genes. (C) *PHI-1* responds to the well-known developmental and environmental cues related to cellulose synthesis. By comparison to the *CESA*s expression, the expression of *PHI-1* was upregulated quite evidently in response to the developmental and environmental signals related to cellulose synthesis. *PHI-1*, AT1g35140; *CESA1*, AT4G32410; *CESA2*, AT4G39350; *CESA3*, AT5G05170; *CESA4*, AT5G44030.

PHI-1 was excluded as a general regulator of the cell cycle owing to its non-cyclic expression pattern during the cell cycle progression of tobacco BY-2 cells. However, its expression could be induced when phosphate-starved tobacco BY-2 cells re-enter cell cycle from a static state in the presence of phosphate (Toshio et al., 1999). Since primary cell wall synthesis is one of the scenarios during plant cell cycle progression (Somerville et al., 2004), the behavior of *PHI-1* in cultured tobacco BY-2 cells led us to hypothesize its role in regulating the synthesis of wall polymers, particularly cellulose, in newly formed cells. Moreover, the fact that its homologous gene was involved in cell morphogenesis of the giant cells induced by the root-knot nematode *Meloidogyne javanica* in tomato further provides potential evidence for the PHI-1 function in the regulation of cellulose biosynthesis (Fosu-Nyarko et al. 2009). In early-stage *M. javanica*-induced giant cells, it was observed that the transcript abundance of its homologous gene was 6.7∼8.5-fold higher than that in control cells. During this stage, the giant cells expand rapidly, reflecting a demand for prompt cellulose biosynthesis, but none of the known cellulose biosynthesis-related genes was upregulated (Fosu-Nyarko et al. 2009).

Since the surface of cellulose microfibrils is coated by cross-linking hemicelluloses and the cellulose deposition in the primary cell wall accompanies cell expansion (Scheller and Ulvskov, 2010), the regulatory component of a CSC, which functions during a primary cell wall formation should co-express with the genes responsible for either hemicellulose synthesis or cell expansion. Further analysis of the genes co-expressed with *PHI-1* by the public database ATTED-II showed that PHI-1 could be such a candidate because two top genes co-expressed with *PHI-1*, *XYLOSYLTRANSFERASE 1* (*XT1*) and *ENDOTRANSGLUCOSYLASE/HYDROLASE 22* (*TCH4*), meet this requirement (Obayashi et al., 2022) (Table 1). XT1 is involved in xyloglucan biosynthetic process, while TCH4 is involved in xyloglucan metabolic process as cell wall-modifying enzymes for the controlled cell wall expansion (Scheller and Ulvskov, 2010).

**Table 1.** Co-expressed genes with *PHI-1* (top 3) in Arabidopsis. Data from public database ATTED-II (Obayashi et al., 2022).

| Locus | Alias | Function | Involved biological process |
| --- | --- | --- | --- |
| *At5g57560* | *TCH4* | xyloglucan<br>endotransglucosylas<br>e/ hydrolase family<br>protein | Cell wall biogenesis and<br>organization, xyloglucan<br>metabolic process, response to<br>mechanical stimulus,<br>brassinosteroid stimulus |
| *At4g08950* | *EXO* | Phosphate-induced<br>1/Exordium-like<br>family protein | response to brassinosteroid<br>stimulus |
| *At3g62720* | *XT1* | xylosyltransferase 1 | xyloglucan biosynthetic<br>process, response to<br>mechanical stimulus |

A variety of environmental or developmental cues can lead to changes in cell expansion and cell wall synthesis (Sun et al., 2010; Wang et al. 2020). To investigate whether *PHI-1* responds to the well-known developmental and environmental stimuli related to cellulose synthesis, the information from the public microarray database Genevestigator was analyzed (https://genevestigator.com/gv/).

Compared with the expression of *CESAs*, the *PHI-1* expression was significantly upregulated in response to some stresses, such as hypoxia and dark treatments (Figure 1C). A common phenomenon observed when plants respond to these stresses is that their aboveground parts, such as petiole, hypocotyl, and stem, rapidly elongate, indicating a fast cellulose synthesis to maintain cell wall thickness. In Arabidopsis, an increased expression of *PHI-1* was also observed when treated with brassinolide, a member of the brassinosteroids (BRs) (Figure 1C). It was believed that the BR hormones have effects on cell wall formation and carbon partitioning, owing to the fact that they promote cell elongation, signifying an enhancement of cellulose biosynthesis to maintain wall thickness (Sun et al., 2010).

In pollen germination, the cell wall of the tube cell rapidly elongates, also implying a prompt cellulose synthesis. Interestingly, the expression of *PHI-1*markedly rises during pollen tube growth (Figure 1C). Moreover, it was reported that *PHI-1* responds to mechanical stimuli (Heyndrickx and Vandepoele 2012). It has long been believed that tension wood, which has a higher proportion of cellulose than normal wood, was attributed to mechanical stimuli (Guerriero et al., 2014).

For these reasons, PHI-1 is presumably involved in signaling processes that integrate developmental or environmental signals with cellulose biosynthesis during cell wall formation.

### The cell-type-specific expression pattern of *PHI-1* coincides closely with a need for cellulose biosynthesis in plant

The expression pattern of *PHI-1* helps us evaluate the biological processes in which it may be involved. In Arabidopsis, however, different experiments showed different expression patterns of its closest paralogous gene *EXO*, resulting in conflicting judgments on the role of EXO. In the promoter trapping experiment, *EXO* expressed in proliferating cells, thereby playing a role in cell division (Farrar et al., 2003), while in expression profile analysis, *EXO* expression was upregulated by BR induction, thereby playing a role in cell expansion (Schröder et al., 2009).

To thoroughly examine the *PHI-1* expression pattern, we observed β-glucuronidase (GUS) activities *in situ* by expressing *GUS* under the control of *PHI-1* promoter in transgenic lines. The results showed that GUS activities were found in all kinds of plant organs (Figures 2A-2E). A prominent feature is that the expression of *PHI-1* is cell-type-specific. GUS expression was virtually observed neither in fully differentiated cells, such as those in matured leaves (Figure 2D), nor in actively dividing cells, such as those in the apical meristem of caudices (Figures 2A and 2F), while in differentiating cells, strong GUS activities were observed, especially in those undergoing either cell expansion such as cells of elongation zone of primary root and cells of young leaves (Figures 2G and 2H), or primary and secondary cell wall thickening such as stoma guard cells and vascular tissue cells (Figures 2I-2K).

**Figure 2.**
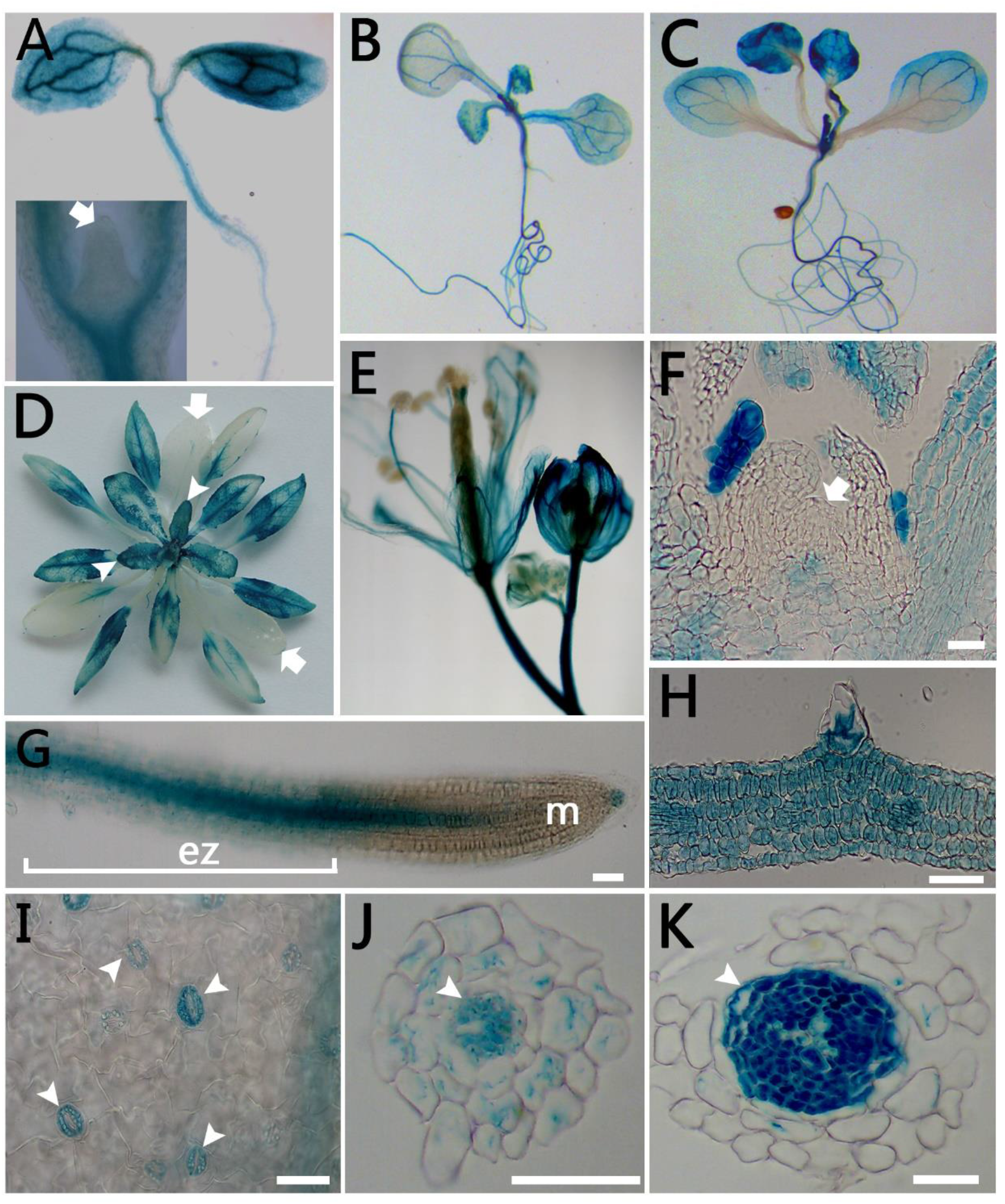
The cell-type-specific expression pattern of *PHI-1* coincides closely with a need for cellulose biosynthesis in plant. (A-E) Surface view of *PHI-1_pro_:GUS* expression throughout development in different organs. (A) Cotyledon stage seedling. The inset is a close-up of the apical meristem (arrow). (B) Two leaves visible stage seedling. (C) Four leaves visible stage seedling. (D) The transition stage from vegetative growth to reproduction. Young leaf (arrowheads); matured leaf (arrows). (E) Flowers and a young silique. (F-K) GUS staining of transgenic plants showing preferential expression in differentiating cells, especially in those undergoing either cell expansion or primary and secondary cell wall thickening. Bar = 30μm. (F) Longitudinal section of a caudex tip. Meristem (arrowhead). (G) Surface view of a root tip. m, meristem; ez, elongation zone. (H) Transverse section of a young leaf blade. (I) Maturing guard cells (arrowheads) of a cotyledon. (J) Transverse section of the distal root hair zone. Vascular tissue (arrowhead). (K) Transverse section of the proximal root hair zone. Vascular tissue (arrowhead).

During cell expansion, an enhancement of cellulose synthesis is required because the wall thickness needs to be maintained (Somerville, 2006). Also, the formation of the lens-thickened wall during guard cell differentiation and the occurrence of secondary wall thickening during vascular differentiation indicate a need for cellulose synthesis. To conclude, the expression pattern of *PHI-1* in Arabidopsis is consistent with a need to synthesize cellulose during cell expansion and cell wall thickening instead of a need for cell division, and coincides closely with the spatial and temporal dependent manner of cellulose synthesis.

### Overexpression of *PHI-1* raised the cellulose content, decreased the cell wall plasticity, and resulted in a distinct alteration of plant architecture

Suppose that PHI-1 is the positive regulatory component of a CSC according to our hypothesis. In that case, its constitutive overexpression will certainly enhance the activities of CSCs, hence leading to a rise in cellulose synthesis. To test our hypothesis, transgenic lines constitutively overexpressing *PHI-1* were generated by expressing *PHI-1* under the control of the *CaMV 35S* promoter in Arabidopsis. In comparison with the wildtype, the *Pro35S:PHI-1* lines presented a little delay of development, and their leaves exhibited a darker green color and their architecture also distinctly altered, displaying a significantly reduced leaf size (Figures 3A and 4E-4H) and a lower height of central stem (Figure 3B).

**Figure 3.**
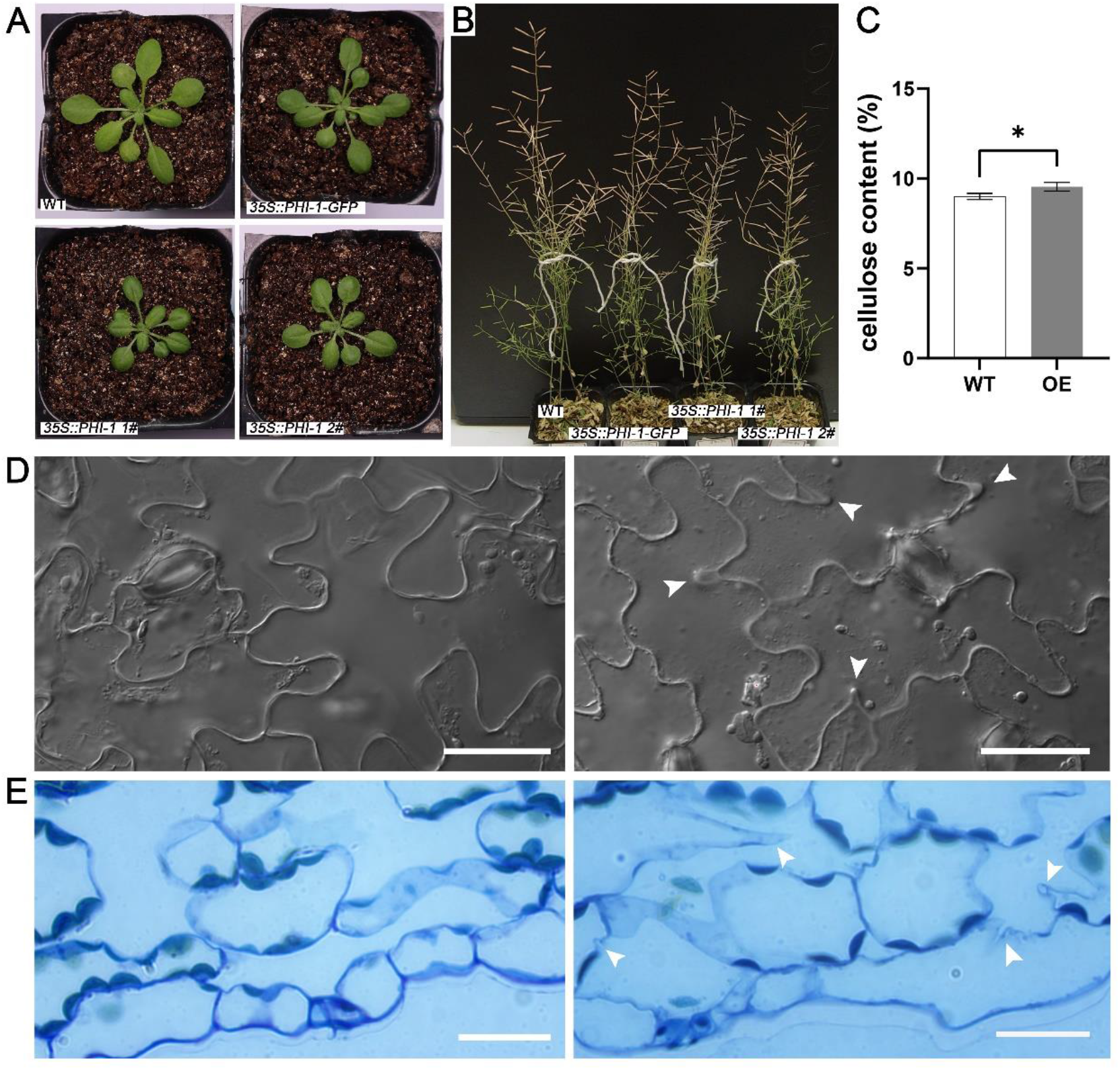
Overexpression of *PHI-1* raised the cellulose content, decreased the cell wall plasticity, and resulted in a distinct alteration of plant architecture. (A) Different plant architectures among the wildtype and *Pro35S:PHI-1*/*Pro35S:PHI-1-GFP* lines. The wildtype and three independent *Pro35S:PHI-1*/*Pro35S:PHI-1-GFP* lines were grown for 21 days. (B) The transgenic lines overexpressing either *PHI-1* or *PHI-1-GFP* presented a lower height of their central stem by comparison with the wildtype, grown for 2 months. (C) The cellulose content of mature rosette leaves was raised in the *PHI-1*-overexpressed line (OE) compared with the wildtype (WT). *, *t* test, significantly different (p<0.05). Error bars represent SD. (D and E) Defects in the control of the anisotropic expansion of cells in the *PHI-1*-overexpressed line. (D) Differential interference contrast (DIC) images of abaxial leaf epidermis. Left, wildtype; Right, *Pro35S:PHI-1* line. Arrowheads indicate more irregular expansion of pavement cell walls of the *PHI-1*-overexpressed line. Bar=30µm. (E) Light microscope images. Left, wildtype; Right, *Pro35S:PHI-1* line. Arrowheads indicate more irregular expansion of the cell wall of mesophyll cells of the *PHI-1*-overexpressed line. Bar=20µm.

**Figure 4.**
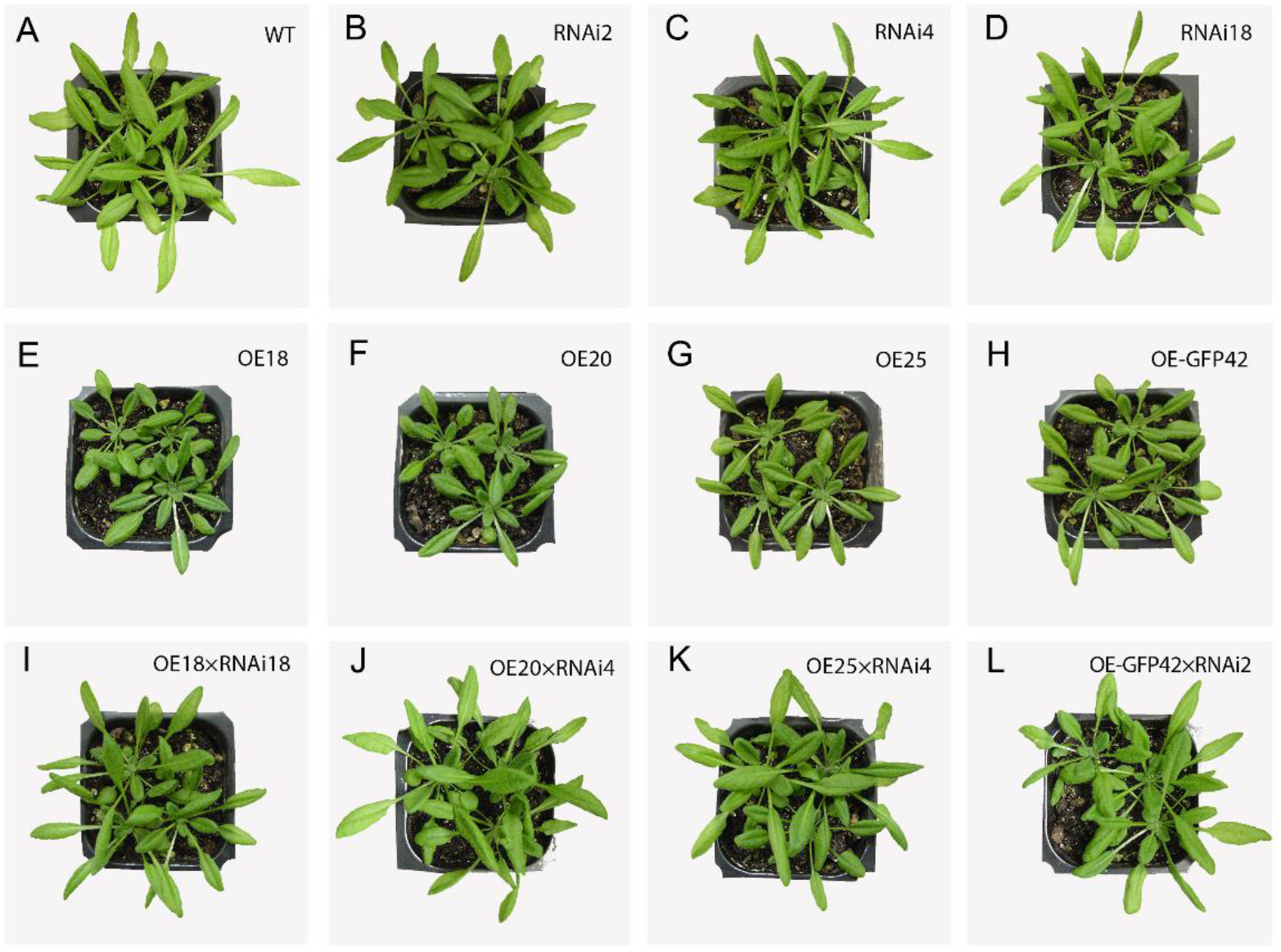
Phenotype analysis of *PHI-1*-overexpressed lines. The hybridization between *PHI-1*/*PHI-1-GFP*-overexpressed lines and *PHI-1* targeted RNAi lines demonstrated that the features of *Pro35S:PHI-1/ Pro35S:PHI-1-GFP* lines truly resulted from the overexpression of *PHI-1*/*PHI-1-GFP*. Plants were grown for 21 days under continuous illumination. (A) The wildtype. (B-D) Plants from different *PHI-1* targeted RNAi lines. Lines # RNAi2, RNAi 4, RNAi 18. (E-G) Plants from different *Pro35S:PHI-1* lines. Lines # OE18, OE20, OE25. (H) A *Pro35S:PHI-1-GFP* plant. Line # OE-GFP42. (I-L) Homozygous hybrids, presenting their phenotype as the wildtype instead of overexpression lines.

This characteristic phenotype of the *Pro35S:PHI-1* lines goes against the common knowledge that cellulose synthesis promotes cell and plant growth (Somerville, 2006), and was not found in *EXO*-overexpressed lines in previous work (Schröder et al., 2009). To further investigate the authenticity of the phenotypic features of *Pro35S:PHI-1* lines, we try to silence the *Pro35S:PHI-1* by RNA interference (RNAi) to observe the morphological change of the *Pro35S:PHI-1* lines. At first, we generated the *PHI-1* targeted RNAi transgenic lines under wildtype background. Each RNAi transgenic line presented a similar phenotype as the wildtype (Figures 4B-4D). After that, we crossed different RNAi lines (carrying a basta-resistant gene) with different *Pro35S:PHI-1* lines (carrying a hygromycin-resistant gene). The phenotype of the hybrids obtained by basta and hygromycin double-resistant screening, harboring both the *Pro35S:PHI-1* and RNAi elements, is in common with that of the wildtype, but did not reflect the features of *Pro35S:PHI-1* lines (Figures 4I-4L). Therefore, RNAi successfully silenced the *Pro35S:PHI-1* in the hybrids and made them exhibit the phenotype of RNAi lines. These results confirmed that the characteristic features of *Pro35S:PHI-1* lines truly resulted from the overexpression of *PHI-1*.

In order to explore the reason why the overexpression of *PHI-1* makes the leaves smaller, we observed the morphology of the leaf epidermis and mesophyll and found that in comparison with those of the wildtype, the leaf epidermis and mesophyll of the *PHI-1*-overexpressed line had noticeable defects in anisotropic cell expansion, manifesting as more irregular expansion of cell wall (Figures 3D and 3E). Meanwhile, we found that the mature rosette leaves of the *Pro35S:PHI-1* lines were highly fragile, and it is much more difficult to peel off their lower leaf epidermis than that of the wildtype, indicating that the cell wall composition could be changed. Then, we measured the crystalline cellulose (acetic-nitric acid-resistant (1, 4) -β-glucan) in dry mature rosette leaves of the *Pro35S:PHI-1* line and found about a 6% increase compared with those of the wildtype (Figure 3C). Thus, we verified that PHI-1 is really a regulatory protein in cellulose biosynthesis.

### PHI-1 and its paralogs possess highly overlapping functions and the mutants of loss-of-function of the entire PHI-1 family presented cellulose deficiency

The mutants of loss-of-function of PHI-1 family proteins will help assess their function precisely *in planta*. A T-DNA insertion mutant of *PHI-1* (*phi-1*) was identified (Supplemental Materials 1). This insertion mutant of PHI-1 showed no visualized change and presented a similar phenotype as the wildtype (Figure 5). EXO is the closest PHI-1 paralog (76% identity of amino acid sequence) (Farrar et al., 2003). *EXO* has an overlapping expression pattern with *PHI-1* (Figure 6A) and the *Pro35S:EXO* line presents the same phenotype as the *Pro35S:PHI-1* line (Figure 6D). We also identified a T-DNA insertion mutant of *EXO* (*exo*) (Supplemental Materials 1). Interestingly, not only the insertion mutant of EXO but also the *exo phi-1* double mutant is similar to the *phi-1*mutant displaying no observable change of morphological characteristics compared with the wildtype (Figure 5). Consequently, we believed that both PHI-1 and EXO are functionally redundant with the other members of the PHI-1 protein family.

**Figure 5.**
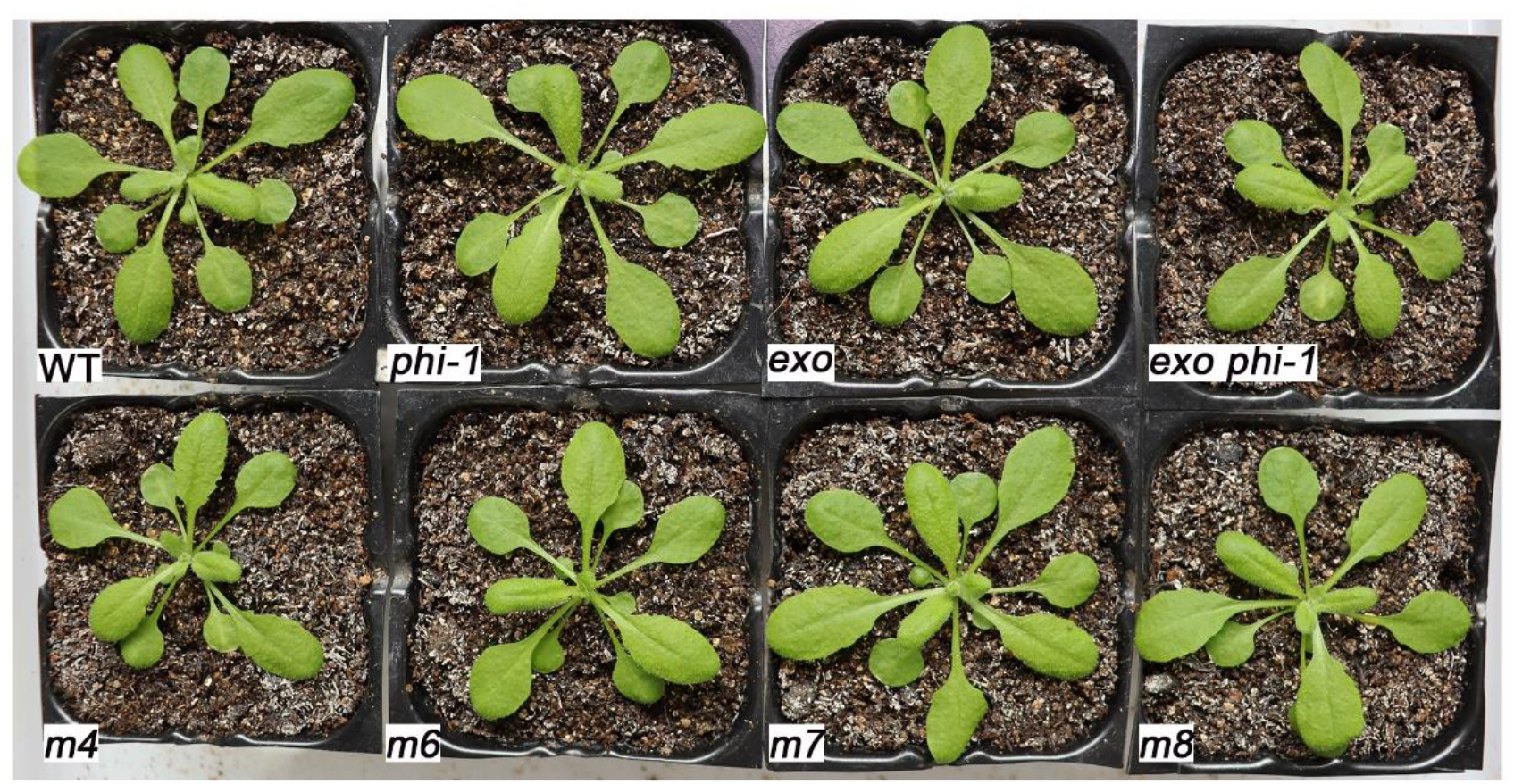
Phenotype analysis of mutants. Different order mutants, *phi-1*, *exo*, *exo phi-1*, *exo exl2 exl4 phi-1*(*m4*), *exo exl2 exl4 exl5 exl6 phi-1*(*m6*), *exo exl2 exl3 exl4 exl5 exl6 phi-1*(*m7*), *exo exl2 exl3 exl4 exl5 exl6 exl7 phi-1*(*m8*), presented no clearly observable change of morphological characteristics during vegetative growth compared with the wildtype (WT) (grown for 21 days).

**Figure 6.**
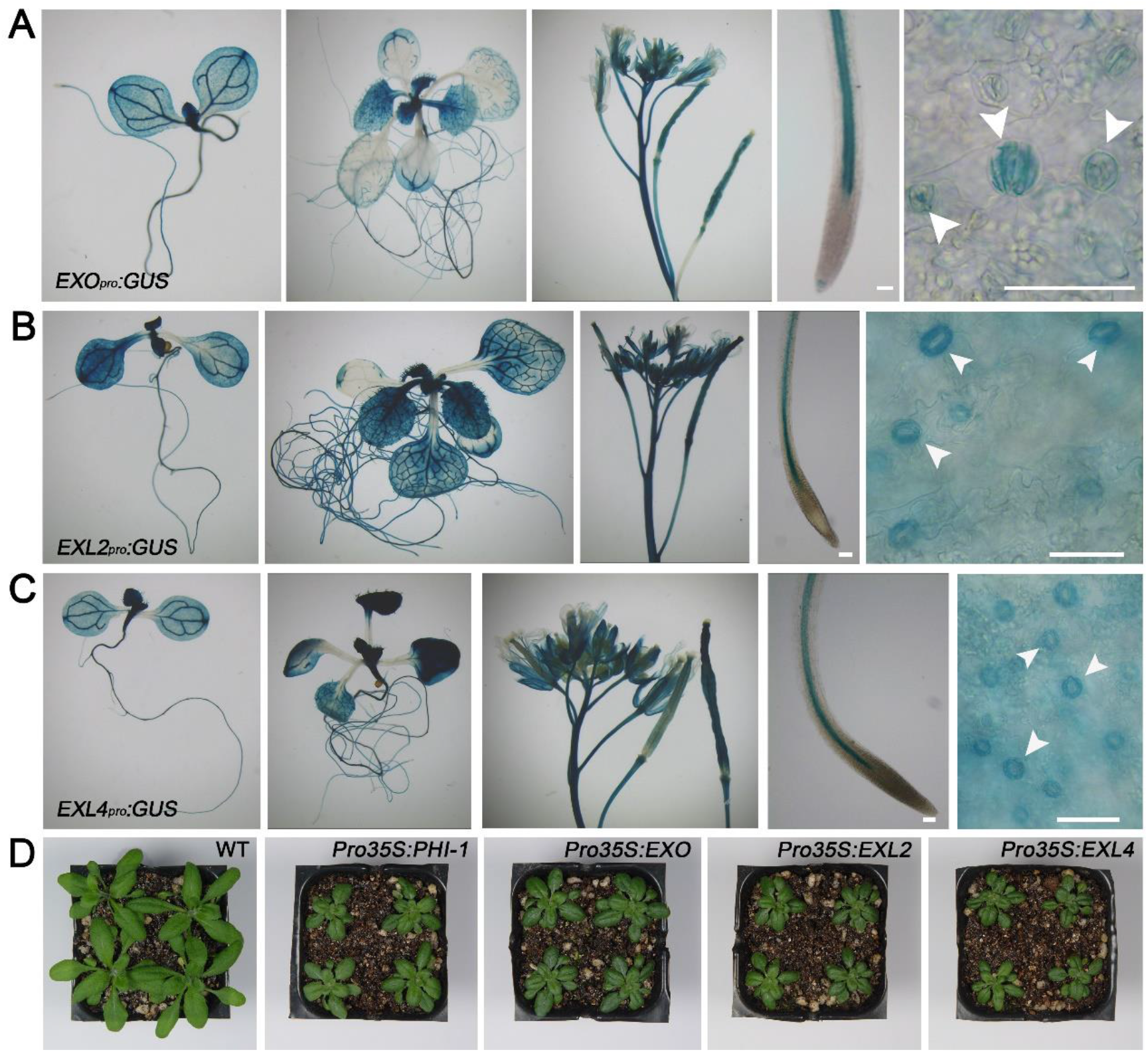
PHI-1 and its paralogs possess highly overlapping functions. (A-C) *EXO* and *EXL2* and *EXL4* all have the overlapping expression patterns with *PHI-1*. GUS staining of transgenic plants showed that *EXO_pro_:GUS* and *EXL2 _pro_:GUS* and *EXL4 _pro_:GUS* all expressed throughout development and exhibited a cell-type-specific pattern, preferentially expressing in differentiating cells, especially in those undergoing either cell expansion or primary and secondary cell wall thickening, just as the characteristic of *PHI-1 _pro_:GUS*. Left to right: two leave visible stage seedlings; six leave visible stage seedlings; flowers and young siliques; surface view of a root tip; and maturing guard cells (arrowheads) of a cotyledon. Bar = 60μm. See also Figure 2. (D) The *Pro35S:EXO* and *Pro35S:EXL2* and *Pro35S:EXL4* lines all present the same phenotype as the *Pro35S:PHI-1* line.

Phylogenetic analyses showed that EXL2 and EXL4 are the next closest related paralogs to PHI-1 (Figure 1A). The *GUS* reporter gene revealed that both *EXL2* and *EXL4* have overlapping expression patterns with *PHI-1* (Figures 6B and 6C), and either *Pro35S:EXL2* or *Pro35S:EXL4* lines present the same phenotype as the *Pro35S:PHI-1* line (Figure 6D). Thus, PHI-1 and its paralogs seemed to possess highly overlapping functions, and the PHI-1 paralogs may compensate for their role in cellulose synthesis in mutants. This encouraged us to establish higher-order mutants to elucidate their relationship with PHI-1.

To probe the function of both PHI-1 and its paralogs further, we generated an *exl2 exl4* double mutant by CRISPR/Cas9 gene editing technology (Supplemental Materials 2). No visible change of morphological characteristics was observed in *exl2 exl4* double mutants in comparison with the wildtype. Then, by self-fertilizing a heterozygote originated in a crossbreeding between an *exo phi-1* double mutant and an *exl2 exl4* double mutant, we obtained an *exo exl2 exl4 phi-1* quadruple mutant among its progenies. Surprisingly, we still failed to characterize any clearly visible phenotypic alteration of the e*xo exl2 exl4 phi-1* quadruple mutant (Figure 5). This means that the members of the PHI-1 family protein function each other in a highly redundant manner in Arabidopsis.

Subsequently, we obtained different mutants via either CRISPR/Cas9 gene editing technology or genetic crossing, including an *exo exl2 exl4 exl5 phi-1* quintuple mutant, an *exo exl2 exl4 exl6 phi-1* quintuple, an *exo exl2 exl4 exl5 exl6 phi-1* sextuple mutant, an *exo exl2 exl3 exl4 exl6 phi-1* sextuple mutant, an *exo exl2 xl3 exl4 exl5 exl6 phi-1* septuple mutant and an *exo exl2 xl3 exl4 exl5 exl6 exl7 phi-1*octuple mutant (Supplemental Materials 2). By comparison with the wildtype, these different higher-order mutants showed no visible morphological changes in vegetative growth (Figure 5), and also exhibited no abnormal phenotype except the *exo exl2 xl3 exl4 exl5 exl6 exl7 phi-1* octuple mutant in reproductive growth. Specifically, the *exo exl2 xl3 exl4 exl5 exl6 exl7 phi-1* octuple mutant has clearly shorted siliques (Figure 7A) and an approximately 23% reduction in seed amount compared with the wildtype (Figure 7B), and its stem also seems more flexible than that of the wildtype (Figure 7C). Then we decided to observe the transverse structure of the stem of the octuple mutant and found that the cells in pith exhibit an irregular cell wall and even broken shapes in comparison with that of the wildtype (Figures 7E and 7F), which is consistent with what is seen in the *cesa1* mutant (Figures 7G and Supplemental Materials 3). The results of cellulose measurement showed that the cellulose content of the main stem of the octuple mutant reduced about 7% compared with the wildtype (Figure 7D). In addition, the result of complementary test with full length *PHI-1* showed that the octuple mutant could be restored as the wildtype phenotype. This means that PHI-1 family members function in a manner of functional overlap and high redundancy, enhancing the control of cellulose biosynthesis when the internal and external environment changes (Kafri et al., 2009).

**Figure 7.**
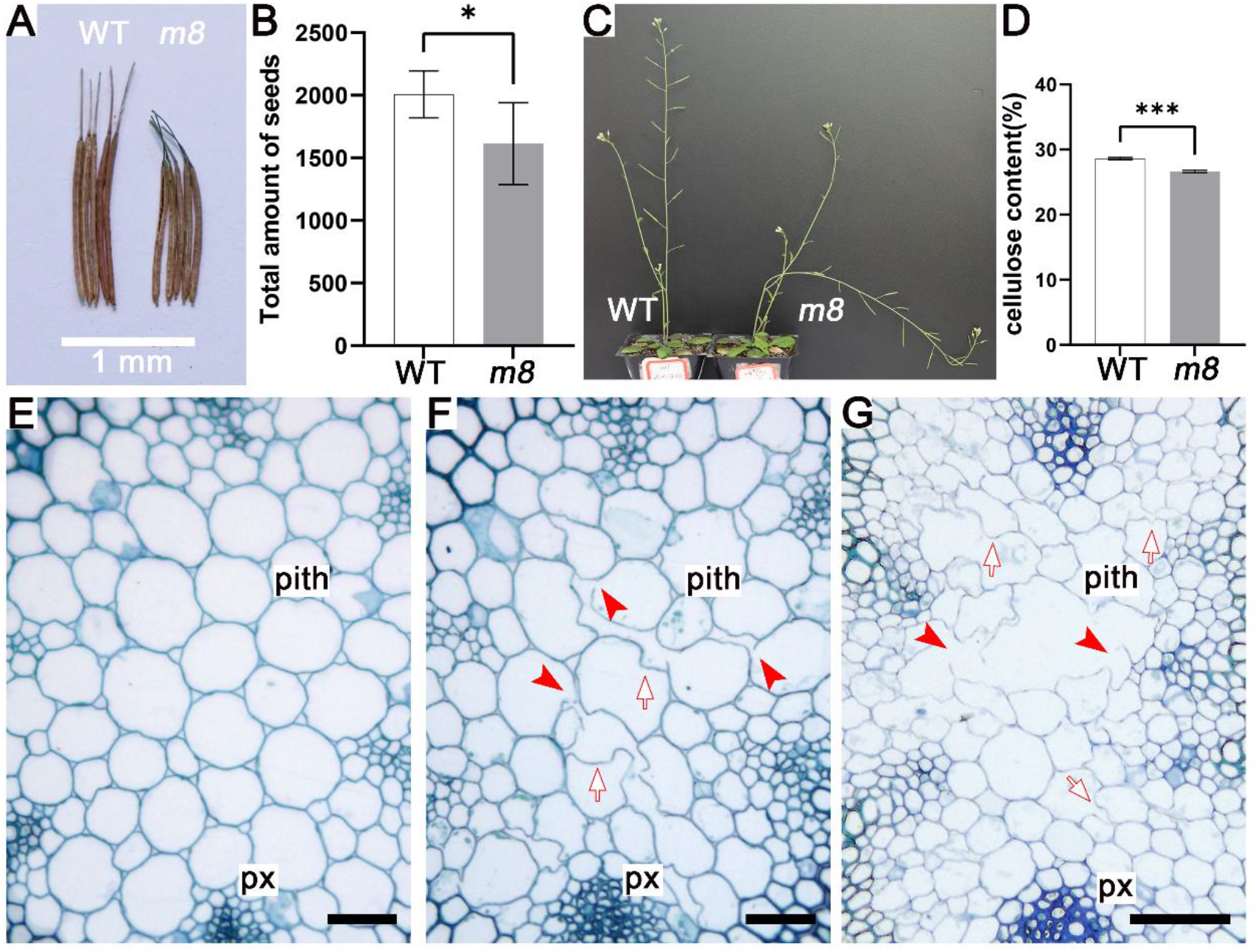
The mutants of loss-of-function of the entire PHI-1 family presented cellulose deficiency. (A and B) The *exo exl2 exl3 exl4 exl5 exl6 exl7 phi-1* octuple mutants of the loss-of-function of the entire PHI-1 protein family presented shorted siliques and had an approximate 23% reduction of seed amount compared with the wildtype. Collected and observed from the sixth to tenth siliques on main inflorescences; six replicates; each group has ten plants at a time. (A) Comparison of silique length. WT, wildtype; *m8*, *exo exl2 exl3 exl4 exl5 exl6 exl7 phi-1* octuple mutant. (B) The number of seeds is reduced about 23% in the octuple mutants (*m8*) compared with the wildtype (WT). *, *t* test, significantly different (p<0.05). Error bars represent SD. (C-G) The loss-of-function of the entire PHI-1 protein family resulted in a cellulose deficiency. (C) The stem of the *exo exl2 exl3 exl4 exl5 exl6 exl7 phi-1* octuple mutant appears more flexible than that of the wildtype. WT, wildtype; *m8*, *exo exl2 exl3 exl4 exl5 exl6 exl7 phi-1* octuple mutant. Grown for 35 days. (D) The cellulose content of the main stems was reduced in the octuple mutant (*m8*) compared with the wildtype (WT). *, *t* test, significantly different (p<0.05). Error bars represent SD. (E-G) Stem cross sections showed that the pith cells of the octuple mutant exhibited weakened cell anisotropy, displaying irregular cell walls and swollen or broken shapes as the *cesa1* mutant. Arrows indicate irregular cell walls; Arrowheads indicate broken cells. px, primary xylem. Bar = 50µm. (E) Wildtype. (F) Octuple mutant. (G) *cesa1* mutant.

### PHI-1 and its paralogs are plasma membrane-resident proteins and PHI-1 can interact directly with multiple CESAs and its overexpression rescued *cesa1*mutant

PHI-1 does not share any significant homology with any protein with known functions over its full length (Toshio et al., 1999). However, PHI-1 shared limited homology with plasma membrane H^+^-ATPases of fungi and plants (Toshio et al., 1999). Thus, it is speculated that PHI-1 may play a certain role in phosphorylation due to the ATP binding site at its N-terminus (Toshio et al., 1999). Furthermore, some works showed that the CSC activities were affected by the phosphorylation of CESAs (Chen et al. 2010, 2016; Bischoff et al., 2011).

The subcellular localization of PHI-1 would contribute to clarifying its molecular function and the biological process involved. In Arabidopsis, its closest paralog EXO was identified as an extracellular protein using fluorescently tagged EXO (Schröder et al., 2009). Proteomic analyses showed that EXL4 (EXORDIUM LIKE 4) of PHI-1 protein family was classified as a membrane protein containing at least one putative transmembrane domain (Mitra et al., 2007), while PHI-1 as well as another two members EXO and EXL2 were classified as potential extracellular proteins because of their signal peptides (Borderies et al., 2003; Bayer et al., 2006). However, the cell wall separation method used in the latter two proteomic experiments cannot rule out a possibility that these cell-wall-associated PHI-1 family proteins may be plasma membrane proteins loosely bound to the cell wall because some transmembrane proteins have a signal peptide just as the cell wall proteins (secreted protein). In addition, the predicted structure *in silico* showed that PHI-1 could be a single-pass transmembrane protein containing a signal peptide at the N terminal (https://phobius.sbc.su.se) (Figure 8A). Therefore, PHI-1 may be a plasma membrane protein closely related to the cell wall.

**Figure 8.**
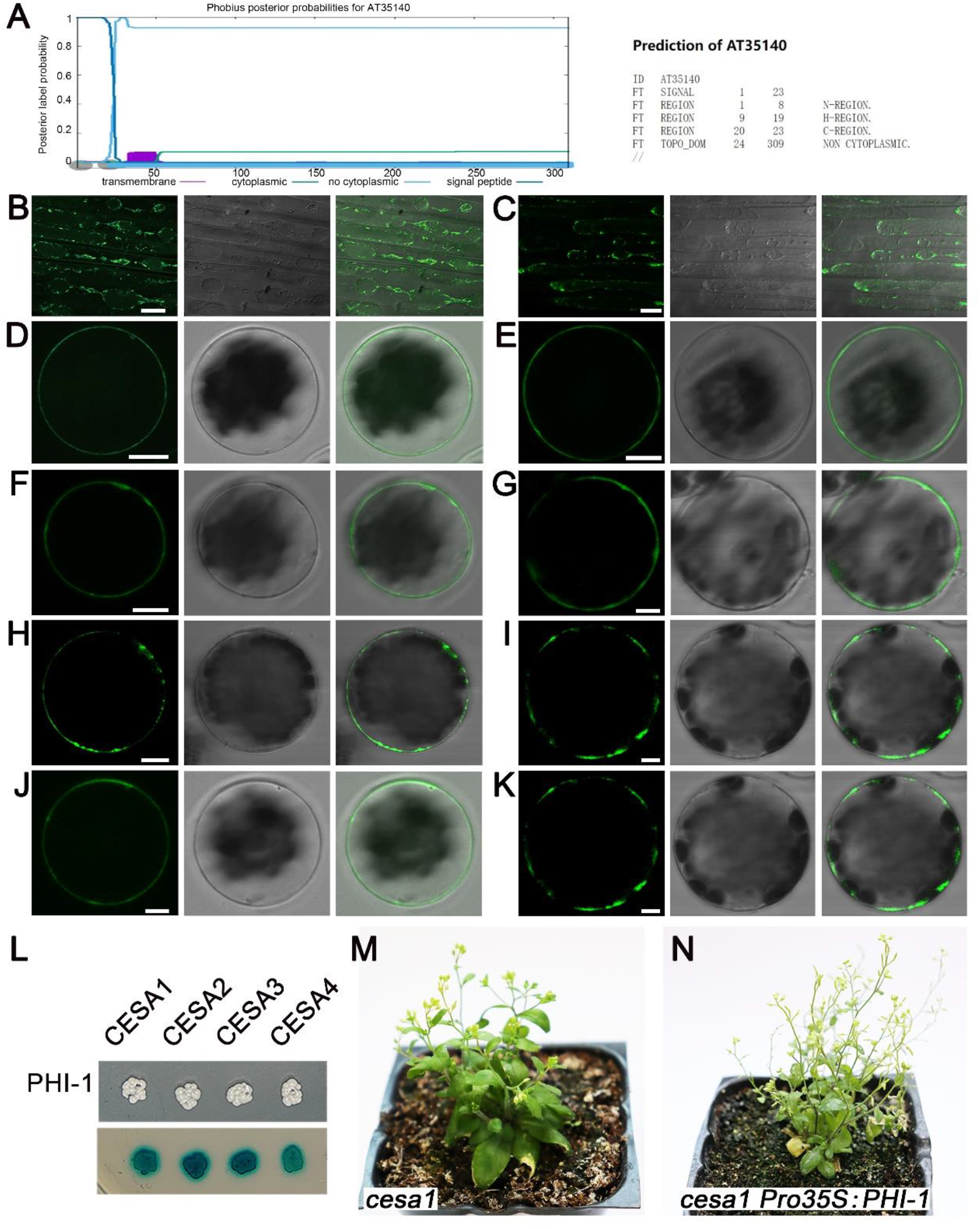
PHI-1 and its paralogs are plasma membrane-resident proteins and PHI-1 can interact directly with multiple CESAs and its overexpression rescued *cesa1*mutant. (A) The predicted structure *in silico* showed that PHI-1 could be a single-pass transmembrane protein containing a signal peptide at the N terminal. The purple column shows the possible transmembrane region. (B-C) Confocal or DIC images of the plasmolyzed hypocotyl cells of a dark-grown either *Pro35S:PHI-1-GFP* or *Pro35S:EXO-GFP* seedling in the presence of 25% (w/v) sucrose. Detection of GFP fluorescence showed that both PHI-1-GFP and EXO-GFP fusion proteins were located in the protoplast. Left to right: a confocal image, a bright-field DIC image and a merged image. Bar = 20μm. (B) PHI-1-GFP. (C) EXO-GFP. (D-K) Confocal or DIC images of a mesophyll protoplast, showing that detection of YFP fluorescence verified that fusion proteins generated by fusing different PHI-1 family proteins with YFP were located at the plasma membrane. The YFP coding sequence was fused to the C terminus of each PHI-1 family protein-coding region in the pA7-YFP vector and then transformed into Arabidopsis mesophyll protoplasts. Left to right: a confocal image, a bright-field DIC image and a merged image. Bar = 10μm. (D) PHI-1-YFP. (E) EXO-YFP. (F) EX2-YFP. (G) EXL3-YFP. (H) EXL4-YFP. (I) EXL5-YFP. (J) EXL6-YFP. (K) EXL7-YFP. (L-N) PHI-1 can interact with multiple CESAs and its overexpression rescued *cesa1*mutant. (L) Interactions between PHI-1 and four CESAs were confirmed by growing diploids on selective plates (SC/-Ade-His-Leu-Trp; white) and an X-α-gal assay (blue) in yeast. (M) A *cesa1*mutant generated by CRISPR/Cas9 gene editing technology, which is infertile, grown for 2 months. (N) A fertile *cesa1*mutant under the *PHI-1*-overexpressed background, grown for 2 months.

For more accurately localizing PHI-1, we constitutively overexpressed PHI-1-GFP fusion protein in Arabidopsis. The *Pro35S:PHI-1-GFP* line presents a similar phenotype as the *35S::PHI-1* line (Figure 3A and 3B), indicating that the C-terminally GFP-tagged PHI-1 possesses an identical function with native PHI-1. Figure 8B shows that GFP-tagged PHI-1 is localized in the protoplast of the plasmolyzed hypocotyl cells of a dark-grown *Pro35S:PHI-1-GFP* seedling. We also substantiated that EXO-GFP is localized in the protoplast instead of in the cell wall in the plasmolyzed hypocotyl cells of a dark-grown *Pro35S:EXO-GFP* seedling (Figure 8C). Furthermore, we transiently expressed PHI-1-YFP fusion protein and other fusion proteins in mesophyll protoplasts, by fusing each PHI-1 family protein with YFP, and observed clearly that each C-terminally YFP-tagged PHI-1 family protein is sited on the plasma membrane (Figures 8D-8K).

Therefore, both PHI-1 and its paralogs are plasma membrane proteins instead of cell wall proteins, and it should be possible that these proteins could play a role in the phosphorylation of CESAs owing to their putative phosphorylation function and the same plasma membrane localization with CESAs.

Since both PHI-1 and CESAs are plasma membrane-resident proteins, their physical interaction is possible. To investigate whether there are interactions between PHI-1 and CESAs, we made use of a mating-based split ubiquitin system of yeast two-hybrid to test membrane protein interactions. PHI-1 was used as a bait protein, and CESAs as prey proteins. Interactions were selected on SC/-Ade-His-Leu-Trp media. Figure 8L showed that PHI-1 could interact at least with four CESA members, including CESA1, CESA2, CESA3, and CESA4, and these interactions were confirmed by an X-α-Gal assay.

Multiple CESAs are involved in the formation of the primary cell wall, among which CESA1 is considered essential (Polko and Kieber 2019). Because PHI-1 can interact with multiple primary wall CESAs, we want to test whether the overexpression of *PHI-1* can rescue the *cesa1* mutant by strengthening the interaction between PHI-1 and the remaining primary cell wall CESAs. By means of CRISPR/Cas9 gene editing technology, we generated a *cesa1* mutant (Figures 8M and S2J). This mutant is infertile, but it can be cross-fertilized with the wildtype pollens, meaning that its male gametophyte malfunctions. Then, we tried to pollinate this *cesa1* mutant with the *Pro35S:PHI-1* line pollens, and directly obtained the fertile *cesa1* mutant under *PHI-1*-overexpressed background due to the continued effect of CRISPR/Cas9 (Figure 8N) and the sequencing result of PCR product for its identification is identical as that of the *cesa1* mutant (Supplemental Figure 2J). This result proves that by enhancing the interaction between PHI-1 and the remaining primary cell wall CESAs, cellulose synthesis in *cesa1* mutants can be improved so that the mutant, benefiting from a functional male gametophyte, can successfully achieve alternation of generations.

The above experimental results suggested that PHI-1 is directly involved in the regulation of cellulose biosynthesis by its interaction with CESAs in Arabidopsis.

## Discussion

Cell walls define the shape of plant cells, controlling the extent and orientation of cell elongation, and hence plant growth. The main load-bearing component of plant cell walls is cellulose, and how plants regulate its biosynthesis during development and in response to various environmental perturbations is a central question in plant biology (Polko and Kieber 2019). In this study, we demonstrated that PHI-1 and its paralogs positively regulate cellulose biosynthesis in Arabidopsis.

### Subcellular localization of PHI-1 family proteins

Protein subcellular localization is very important for studying protein function, which provides a research direction for understanding the mechanism of action of a protein, such as the possible place where it can function and its interaction with proteins at the same subcellular location. Our findings demonstrated that PHI-1 family proteins are plasma membrane-resident instead of cell wall-resident as reported before.

There are several reasons why the previous work failed to make the correct subcellular localization of PHI-1 family proteins (Borderies et al., 2003; Bayer et al., 2006; Schröder et al., 2009). Firstly, PHI-1 family proteins have no homology with any other proteins with known functions (Toshio et al., 1999); secondly, both structure analyses of PHI-1protein family *in silico* and proteomic analyses classified preferentially PHI-1 family members as secreted proteins due to their signal peptides at the N terminal, although PHI-1 has clearly a predicted transmembrane hydrophobic domain (Figure 8A); moreover, plant cell walls tend to produce strong autofluorescence, meanwhile the fusion reporter gene localization method makes the fluorescent signal of the fusion protein scattered on the plasma membrane much weaker than that concentrated in organelles such as the nucleus, and especially, the expression level of *Pro35S:EXO-GFP* were not high enough to be detected owing to the selected line did not show the characteristic phenotype identified in this study in PHI-1 family gene-overexpressed lines (Schröder et al., 2009).

### Identification of *PHI-1*-overexpressed lines

The plant cell wall is a dynamic structure that changes in response to developmental and environmental cues, meaning that there is a mechanism to regulate cellulose biosynthesis to adapt to growth and environmental changes (Somerville et al., 2004; Sun et al., 2010; Wang et al. 2020). *CESA* genes appear to be expressed semi-constitutively under developmental programs, but do not respond to environmental stimuli (Somerville, 2006). Therefore, cellulose biosynthesis is not solely controlled at *CESA* transcription level. On the other hand, it was suggested that the CSC activity could be controlled at the posttranscriptional level in plants by a plasma membrane protein interacting with CESA proteins (Somerville et al., 2004; Somerville, 2006; Endler and Persson, 2011).

In this study, we found that constitutive overexpression of *PHI-1* raised the cellulose content, and further experiments explicitly demonstrated that PHI-1 is really a positive regulatory protein in cellulose biosynthesis. However, excessive PHI-1 had a significant impact on the overall architecture of plants, in which the anisotropic cell expansion showed obvious defects (Figures 3D and 3E).

These *PHI-1*-overexpressed plants displaying a defective cell expansion might set up a counterintuitive paradox. Logically, cells require PHI-1 for their normal expansion, due to the fact that *PHI-1* has a very high expression level in rapidly expanding cells (Figures 2G and 2H). Hence, the inhibitive effect of excessive PHI-1 on anisotropic cell expansion must be indirect. Since the xyloglucan endotransglycosylase, encoded by *TCH4*, which co-express most significantly with *PHI-1*, plays a part in controlled cell wall expansion by catalyzing transglycosylation of xyloglucan with molecules bound to cellulose (Chanliaud et al., 2004; Fry, 2004), this defect in anisotropic cell expansion should attribute to a decrease of cell wall plasticity, derived from an asynchronization between cellulose biosynthesis and transglycosylation of xyloglucan with molecules bound to cellulose in the *Pro35S:PHI-1* lines. This experimental result reflected that a well-functioning cell wall resulted from a coordinated expression of multiple associated genes.

### Activation of the inactive CSCs

In plants, the inactive CSCs are assembled in the Golgi apparatus, and moved to the plasma membrane, where they become activated (Haigler and Brown 1986). As yet, we know nothing about the activation mechanism, but it was speculated that the CSCs may be activated via transiently binding an accessory protein, in response to developmental or environmental signals (Somerville, 2006; Endler and Persson, 2011).

In the beginning, we also imagined the role of PHI-1 in activating the inactive CSCs that have just moved to the plasma membrane, because the expression of *PHI-1* responds to the well-known developmental and environmental cues related to cellulose synthesis. However, the *exo exl2 xl3 exl4 exl5 exl6 exl7 phi-1* octuple mutants of the loss-of-function of the entire PHI-1 family are viable. Thus, PHI-1 family proteins are certainly not indispensable for the activation of the inactive CSCs. Other unknown proteins or other unknown mechanisms for the activation of these inactive CSCs should exist.

### The possibility of an alternative signal pathway to replace the PHI-1 protein family

Cellulose-deficient mutants *cesa1* presented sterility. At the same time, this mutant was also manifested in growth retardation and dwarfing, as cellulose deficiency may trigger some feedback mechanisms that slow down plant growth and make smaller plants adapt to the decline of cellulose biosynthesis ability (Figure 8M).

By comparison, the *exo exl2 xl3 exl4 exl5 exl6 exl7 phi-1* octuple mutants of the loss-of-function of the entire PHI-1 protein family merely showed a mild decline in fertility and a more flexible stem, but the overall plant architecture and growth were similar to that of the wildtype. Only those cells with particularly thin cell walls showed weakened anisotropic expansion or rupture of the cell wall (Figure 7F). These mild phenotypic alterations of the octuple mutant may indicate that the PHI-1 family proteins play a fine-tuning role in integrating developmental and environmental signals into cellulose biosynthesis. Alternatively, there are other signaling pathways to replace the role of the PHI-1 protein family in part because multiple pathways control plant growth and development, and no evidence indicates that *PHI-1* and its paralogous genes respond to gibberellins.

### How does PHI-1 family protein mediate the CSC activity

The PHI-1 family proteins do not show similarities to proteins with known functions (Toshio et al., 1999), and thus may have enzymatic or signaling functions that are unknown to date. Much evidence has shown that not only *PHI-1* but also the other members of the *PHI-1* gene family responded to developmental and environmental signals related to cellulose synthesis, such as *EXO* in promoting stem elongation (Yin et al., 2002), and *EXL6* in pollen tube growth (Wang et al., 2008). In particular, EXO was demonstrated as the terminal of the promoting growth hormone BR signaling pathway (Schröder et al., 2009), which fully fits the scenario under which EXO promotes entire CSC activity to increase cellulose synthesis by feeding BR signaling into CSC. By comparison, it is unlikely that BIN2, as a midway component of the BR signaling pathway, negatively regulates cellulose synthesis by directly phosphorylating CESA1 (Sánchez-Rodríguez et al., 2017).

At the molecular level, protein phosphorylation is the most common event during signal transduction, and post-translational phosphorylation of CESAs has been proposed as one mechanism to mediate the dynamic regulation of cellulose biosynthesis in response to a variety of developmental and environmental stimuli (Speicher et al., 2018). The *PHI-1* gene was first isolated because it was rapidly expressed after the addition of phosphate to phosphate-starved tobacco BY-2 cells and acted to cause phosphate-starved cells to re-enter the cell cycle (Toshio et al., 1999). According to its limited homology with plasma membrane H^+^-ATPases of fungi and plants, PHI-1 is presumed to have an ATP-binding site (a conserved sequence Lys-Gly-Ala) at its N-terminal region, and thus it may have some roles in phosphorylation (Toshio et al., 1999). Moreover, Wilson et al. reported that the phosphate-induced cell cycle re-entry of tobacco BY-2 cells showed the transient activation of a MAP kinase activity by re-feeding of phosphate (Wilson et al. 1998). This observation also reminds us of the role of PHI-1 as a kinase to feed developmental and environmental signals directly into CSC activities by phosphorylating CESAs.

In short, our findings reveal a mechanism in which PHI-1 family proteins, as positive regulatory components of the CSC, can dynamically modulate cellulose biosynthesis in response to various developmental and environmental signals. Hence, we have the opportunity to design new cellulosic materials with diverse economic applications by modifying the cellulose ratio in the cell wall.

## Acknowledgments

This work was supported by the National Natural Science Foundation of China (Project Nos. 30270086, 31471200 and J1210026) and the Fundamental Research Funds for the Central Universities (020814380149).

## Author Contributions

F.Y., S.L., and Q.W. designed the experiments. F.Y., S.L., Q.W. and X. W. wrote the paper. F.Y., Q. W., J. Q., Y. L., X. W., T. C., J. Z., S. Y., Z. Z., X. C. and H.J. conducted the experiments.

## Methods

### Plant Materials and Growth Conditions

*Arabidopsis thaliana* ecotypes Colombia (Col-0) plants were grown either in pots containing a 1:3:1 mix of peat: vermiculite: perlite (v/v) or aseptically on Murashige and Skoog (MS) plates (1/2 ×MS salts, 0.8% agar, 1% sucrose, pH 5.7) in petri dishes. For soil-grown plants, seeds were surface sterilized using 75% ethanol, stratified at 4°C for 3 d, and germinated on MS plates and then were transferred to pots. In selection of transgenic plants, seeds were surface sterilized with 75% ethanol for 10 min, stratified at 4°C for 3 d, and germinated on MS plates containing hygromycin or kanamycin sulfate or sulfonamide (40µg/mL) for hygromycin or kanamycin or sulfonamide resistance, or seeds were germinated in pots, treated with 0.01 % Basta containing 0.1% Tween 20 for Basta resistance. Plants were grown in growth cabinets at 22°C under a 16-h-light / 8-h-dark cycle unless indicated as continuous illumination.

### Generation of Transgenic Plants

For generating *PHI-1_Pro_:GUS*, *EXO _Pro_:GUS*, *EXL2 _Pro_:GUS* and *EXL4 _Pro_:GUS*, a genomic DNA fragment (∼1.5kb) upstream from the start codon (ATG) of *PHI-1*, *EXO*, *EXL2* and *EXL4* was cloned into pCAMBIA1301 using *Bam*H I/*Nco* I or *Eco*R I/*Nco* I, respectively. For generating *Pro35S:PHI-1*, the full-length coding sequence of *PHI-1* modified by a same sense point-mutation at #228 bp (C→T) was cloned into pCAMBIA1301 using *Nco* I/ *Eco*O65 I. For silencing *PHI-1* in Arabidopsis with RNA interference technology, a chimeric gene was constructed by adding sense and antisense parts of *PHI-1* coding sequence GTTGCTTCCCTCTCGTCTTCCCGGAGATCGACCATGGCTCAAAATCCCTCAGTCGCCACGTGGTGGAAGACGGTGGAGAAGTATTACCAATTCCGCAAGATGACCACGACACGTGGACTCAGTCTCTCCCTCGGAGAACAGATCCTCGACCAAGGATACTCAATGGGAAAATCTTTAACAGAGAAAAACCTCAAAGACTTGGCCGCAAAAGGTGGCCAAAGCTACGCGGTTAACGTCGTGTTGACCTCAGCTGACGTGACGGTCCAAGGCTTTTGCATGAACAGATGCGGGTCACACGGGACTGGTTCCGGGTCAGGCAAGAAAGGATCAA, into left (*Nco* I**/***Asc* I) and right arm (*Bam*H I/*Sma* I) of pFGC5941’s CHSA intron respectively. For generating *Pro35S:EXO* and *Pro 35S:EXL2* and *Pro 35S:EXL4*, the full-length *EXO* or *EXL2* or *EXL4* coding sequence was respectively cloned into pCAMBIA1301 using *Nco* I/ *Eco*O65 I or into pBI121-eGFP using *Xba* I/*Sac* I. For generating *Pro35S:PHI-1-GFP* and *Pro35S:EXO-GFP*, the full-length *PHI-1*or *EXO* coding sequence was respectively cloned into pBI121-eGFP using *Xba* I/*Bam*H I. For generating the complementary experiment, the full-length *PHI-1* (∼2.5kb, promoter and coding sequence together) was cloned into pCAMBIA1301 using *Eco*R I/*Eco*O65 I. The verified constructs were then transformed into Arabidopsis using Agrobacterium (GV3101)-mediated floral dipping (Clough and Bent 1998). In each transformation, more than 30 transgenic lines (T0 generation) were isolated by hygromycin, kanamycin or Basta resistance respectively. Those showing a Mendelian segregation of the selective resistance among T1 transgenic plants were selected for obtaining a homozygote by self-pollinating. Analyses were performed in homozygous plants of either T2 or T3 generation. The results represented three independent lines exhibiting identical phenotype. In addition, RT-qPCR was used to verify the expression level of *PHI-1* in the *Pro35S:PHI-1* lines and the *PHI-1* targeted RNAi transgenic lines (Supplemental Figure 1).

### GUS Staining

Transgenic plants were stained for β-glucuronidase (GUS) activity essentially according to the procedure described by Jefferson et al. with some modifications (Jefferson et al., 1987). Briefly, plants were fixed in 90% cool acetone on ice for 30 min. After the plants were washed in staining buffer (a solution containing 50 mM phosphate buffer, pH7.2, 0.02% Triton X-100, 2 mM potassium ferricyanide and ferrocyanide), they were submerged and vacuum infiltrated in the staining buffer containing 2 mM 5-bromo-4-chloro-3-indolyl-β-D-glucuronide (X-gluc) for half an hour on ice. Then, samples were left at 37°C overnight. After the reaction was stopped with water, the plants were washed with 70% ethanol and shined several days for bleaching. Specimens were mounted in 50% glycerol and photographed under a light or stereo microscope. Samples for sectioning were dewatered in a gradient of tert-butanol and embedded in paraffin, cut into 6-8 μm, photographed under a light microscope. For different transgenic plants, at least three independent lines presented identical GUS staining results and one of them was exhibited as a representative.

### RNA Extraction and RT-PCR/ RT-qPCR

Total RNA was extracted from 14-d-old, light-grown seedlings using the RNAiso plus reagent (TaKaRa). First strands were synthesized using PrimeScript^®^ 1^st^ Strand cDNA Synthesis Kit (TaKaRa). RT-PCR was used to test for the presence of transcripts. The quantitative RT-PCR was performed on a Thermal Cycler Dice® TP800 (TaKaRa) with the SYBER^®^ Premix *Ex Taq* ^TM^ II kit (TaKaRa) according to the manufacturer’s protocol. Each reaction was performed on a 1:20 dilution of the first cDNA strands, synthesized as described above, in a total reaction of 20 µL. With this dilution, the SYBR green signal was linear. The expression of *actin2* was quantified as a control. The experiment was performed three times on independent biological samples.

### Leaf Epidermis Observation

In order to easily observe with a laser scanning confocal microscope (Olympus FV1000), we used epidermal peels from the lower leaf epidermis of the 7th mature rosette leaves of flowering stage plants and mounted them with 50% glycerol for observation.

### Anatomical Structure Observation of Leaf and Stem

Leaf blade samples from the 7th mature rosette leaves of fruiting stage plants or the middle of the main stem during the flowering stage were fixed in 2.5% (v/v) glutaraldehyde in 0.1M PBS (pH7.2). After being washed in PBS, samples were postfixed in 2% (v/v) OsO_4_ for 1 h and then dehydrated through a gradient of acetone, and embedded in resin (PON 812, SPI, USA). 2.0 μm thin sections were cut. Then, thin sections were stained with 0.2% Toluidine Blue O for observation under a light microscope.

### Crystalline Cellulose Measurement

Crystalline cellulose (acetic-nitric acid-resistant (1, 4) -β-glucan) was measured from mature rosette leaf fixed by 75% ethanol or from mature main stem dried at 105℃ using the Updegraff method (Updegraff, 1969). Data were collected from three technical replicates for each sample.

### The Subcellular Localization of both PHI-1-GFP and EXO-GFP in Transgenic Plant Cells

The seeds were sowed in pots, stratified at 4°C for 3 d; then placed in light for 1 d at 22°C. After that, these seedlings were transferred in darkness for 3 d at 22°C.

The hypocotyl cells of 4-d-old either *Pro35S:PHI-1-GFP* or *Pro35S:EXO-GFP* etiolated seedlings, which can avoid the influence of autofluorescence usually originated from the cell wall, were used for subcellular localization of either PHI-1-GFP or EXO-GFP in the presence of 25% (w/v) sucrose, respectively. Both PHI-1-GFP and EXO-GFP were excited with a 488 nm laser and their fluorescence were observed with a laser scanning confocal microscope (Olympus FV1000).

### The Subcellular Localization of PHI-1-YFP, EXO-YFP, EXL2-YFP, EXL3-YFP, EXL4-YFP, EXL5-YFP, EXL6-YFP and EXL7-YFP in Mesophyll

#### Protoplasts

Transient expression of PHI-1-YFP, EXO-YFP, EXL2-YFP, EXL3-YFP, EXL4-YFP, EXL5-YFP, EXL6-YFP and EXL7-YFP fusion proteins was performed in mesophyll protoplasts for their subcellular localization (Yoo et al., 2007). The YFP coding sequence was fused to the C terminus of each PHI-1 family protein-coding region in the pA7-YFP (*Bam*H I or *Spe* I) and then transformed into Arabidopsis mesophyll protoplasts. PHI-1-YFP, EXO-YFP, EXL2-YFP, EXL3-YFP, EXL4-YFP, EXL5-YFP, EXL6-YFP and EXL7-YFP were excited with a 514 nm laser and their fluorescence were observed with a laser scanning confocal microscope (Olympus FV1000).

#### Yeast Two-Hybrid Assay

The yeast two-hybrid assays were carried out using a mating-based split ubiquitin system (mbSUS) for membrane protein interactions as described “mbSUS tests protocol” (Clontech) (Obrdlik et al., 2004). Both *PHI-1* ORF and *CESA*s ORFs were amplified from first-strand DNA with PrimeScriptRTase (TaKaRa) using primers indicated in Supplemental information 4. To test the interaction between PHI-1 and CESAs, we cloned the prey bait *PHI-1* ORF into pMetYCgate and the prey *CESA*s ORFs into pNXgate33-3HA by co-transforming an appropriate linear vector restricted with *Pst* I/*Hind* III or with *Eco*R I/*Sma* I and the corresponding PCR fragment insert into yeast strains either THY.AP4 or THY.AP5. Positive interactions were detected both via the ability of diploids to grow on SC/-Ade-His-Leu-Trp media and via LacZ-activity by an X-Gal overlay assay.

To verify detected interactions, we isolated the plasmid DNA from positive diploid yeasts and amplified them in *E. coli*. Then, we isolated the plasmid DNA from *E. coli* and verified the expression cassette by sequencing and repeated again this yeast two hybrid assay by using the verified plasmid DNA instead of the corresponding PCR fragment inserts and linear vectors for mbSUS interaction tests as described “mbSUS tests protocol” (Clontech).

#### Mutants

Both *exo* and *phi-1* were T-DNA insertion mutants, and their seeds were purchased from the Arabidopsis Biological Resource Center (Ohio State University, Columbus, OH). An *exo phi-1*double mutant was obtained from a crossing between an *exo* mutant and a *phi-1*mutant. All T-DNA insertion mutants were identified by PCR (Supplemental Materials 1).

An *exl2 exl4* double mutant was constructed by CRISPR/ Cas9-mediated gene editing technology using a binary vector VK005-3 (ViewSolid Biotech, Beijing). An *exo exl2 exl4 phi-1*quadruple mutant was obtained by a genetic crossing between an *exl2 exl4* double mutant and an *exo phi-1*double mutant (Supplemental Materials 2).

Both *exo exl2 exl4 exl5 phi-1*quintuple mutant and *exo exl2 exl4 exl6 phi-1* quintuple mutant were constructed in the *exo exl2 exl4 phi-1* quadruple mutant background by CRISPR/ Cas9-mediated gene editing technology using a binary vector pKSE401 (a gift from Qi-Jun Chen), respectively. An *exo exl2 exl4 exl5 exl6 phi-1* sextuple mutant was obtained by a genetic crossing between an *exo exl2 exl4 exl5 phi-1* quintuple mutant and an *exo exl2 exl4 exl6 phi-1* quintuple mutant (Supplemental Materials 2).

An *exo exl2 exl3 exl4 exl6 phi-1* sextuple mutant was constructed in the *exo exl2 exl4 exl6 phi-1* quintuple mutant background by CRISPR/ Cas9-mediated gene editing technology using the binary vector VK005-3. An *exo exl2 exl3 exl4 exl5 exl6 phi-1* septuple mutant was obtained by a genetic crossing between an *exo exl2 exl4 exl5 phi-1* quintuple mutant with the *exo exl2 exl3 exl4 exl6 phi-1* sextuple mutant (Supplemental Materials 2).

An *exo exl2 xl3 exl4 exl5 exl6 exl7 phi-1* octuple mutant were obtained by further mutating both the coding region of *PHI-1* and *EXL7* in the *exo phi-1 exl2 exl3 exl4 exl5 exl6* septuple mutant background by CRISPR/ Cas9-mediated gene editing technology using the binary vector VK005-3 (Supplemental Materials 2).

A *cesa1* mutant was constructed by CRISPR/ Cas9-mediated gene editing technology using the binary vector pKSE401 (Supplemental Materials 3).

#### Phylogenetic Analysis

The amino acid sequences were obtained from the Arabidopsis Information Resource and aligned with MUSCLE (Edgar, 2004). The evolutionary history was inferred by using the Maximum Likelihood method based on the JTT matrix-based model (Jones et al., 1992). Evolutionary analyses were conducted in MEGA7 (Kumar et al., 2016).

#### Accession Numbers

*PHI-1*, AT1g35140; *EXO*, AT4G08950; *EXL2*, AT5G64260; *EXL3*, AT5G51550; *EXL4*, AT5G09440; *EXL5*, AT2G17230; *EXL6*, AT3G02970; *EXL7*, AT2G35150; *CESA1*, AT4G32410; *CESA2*, AT4G39350; *CESA3*, AT5G05170; *CESA4*, AT5G44030; *XT1*, AT3G62720; *TCH4*, AT5G57560; *XTR6* AT4G25810; *EXLA1*, AT3G45970.

## Supplemental Materials

1. Identification of the T-DNA insertion mutants (*phi-1* and *exo*) and the establishment and identification of an *exo phi-1* double mutant.
2. Establishment and identification of various mutants targeted to the PHI-1 family by CRISPR/ Cas9-mediated gene editing technology (CRISPR/ Cas9).
3. Establishment and identification of the *cesa1* mutant by CRISPR/ Cas9.
4. List of PCR Primers in this study.
5. Supplemental figure legends.

**Supplemental Figure 1.** Analysis of *PHI-1*expression level in a representative of *35S::PHI-1* lines (OE) and in a representative of *PHI-1* targeted RNAi lines (RNAi) compared to the wildtype (WT) by RT-qPCR. The level of *PHI-1* expression is more than 30 times in the *35S::PHI-1* line and less than 5% in the RNAi line respectively compared with the wildtype, indicating that the expression level of *PHI-1* was just desired in the *35S::PHI-1* line or in the RNAi line. *, *t* test, significantly different (p<0.05). Error bars represent SD. Related to Figure 3 and Figure 4.

**Supplemental Figure 2.** The sequencing result of PCR product for identifcating different mutants. Wildtype (WT) sequence was obtained from TAIR and aligned with BLASTN 2.9.0+ (Altschul et al., 1997). Sequencing peaks presented by Chromas. (A and B) Identification of the T-DNA insertion mutants (*phi* and *exo*). (C-I) Identification of the seven mutated genes of the *PHI-1* family generated by CRISPR/Cas9. Arrows indicated either the base insertion sites or the base deletion site. (J) Identification of *cesa1* generated by CRISPR/Cas9. The arrow indicated a T insertion site. Related to Figure 5 and Figure 7 and Figure 8.

